# Identifying the Species Threat Hotspots from Global Supply Chains

**DOI:** 10.1101/076869

**Authors:** Daniel Moran, Keiichiro Kanemoto

**Affiliations:** Norwegian University of Science and Technology, Trondheim, Norway; Shinshu University, Matsumoto, Japan

## Abstract

**Summary Sentence:** Spatially explicit footprints make it possible to locate biodiversity hotspots linked to global supply chains.

Identifying species threat hotspots has been a successful approach for setting conservation priorities. One major challenge in conservation is that in many hotspots export industries continue to drive overexploitation. Conservation measures must consider not just the point of impact, but also the consumer demand that ultimately drives resource use. To understand which species threat hotspots are driven by which consumers, we have developed a new approach to link a set of biodiversity footprint accounts to the hotspots of threatened species on the IUCN Red List. The result is a map connecting global supply chains to impact locations. Connecting consumption to spatially explicit hotspots driven by production has not been done before on a global scale. Locating biodiversity threat hotspots driven by consumption of goods and services can help connect conservationists, consumers, companies, and governments in order to better target conservation actions.

## Introduction

Human-induced biodiversity threats, such as from deforestation, overfishing, overhunting, and climate change, often arise from incursion into natural ecosystems in search of food, fibre, and resources. A major driver of this incursion is the production of goods for export. Lenzen and colleagues suggested that at least one third of biodiversity threats worldwide are linked to production for international trade^1,2^. Understanding market forces and using effective spatial targeting are key to implementing protections efficiently^3,4^. However, in order to realistically expedite remedial actions threat causes must be located more specifically. Previous work has linked consumption and supply chains to biodiversity impacts but only at the country level^1^. Biodiversity threats are often highly localized. Knowing that a given consumption demand drives biodiversity threat somewhere within a country is not enough information to act. Here we present a new approach to making the inshore and terrestrial biodiversity footprint spatially explicit at a sub-national level.

## Methods

Using a threat hotspot map built using composited extent-of-occurrence maps from IUCN^5^ and BirdLife International^6^ (also called distribution maps showing range boundaries), we applied the biodiversity footprint method of Lenzen *et al*^1^, attributing each anthropogenic species threat to one or more culpable industries then traced the implicated commodities from 15,000 production industries worldwide to final consumers in 187 countries using a global trade model^7,8^. The result is an account linking production and consumption of economic sectors to spatially explicit species threat hotspots. The account only considers threats which can be attributed to industries and thus excludes threats such as change in population structure, disease, natural catastrophes, etc. (discussed further below).

Figure 1 provides an illustration of the method. Species come under threat from a variety of causes, many of which are anthropogenic and linked to industries producing goods for consumption domestically and abroad. In Fig. 1 the extent of occurrence (EOO) of *Ateles paniscus*, the Red-faced Spider Monkey in Brazil, is shaded in a darker colour reflecting that a higher fraction (2.43%) of the total anthropogenic threat to the species that can be attributed to consumption in the US (via agriculture and logging activity in Brazil producing goods finally consumed in the USA). The range of *Atelopus spumarius*, Pebas stubfooted toad, is shown in lighter colour since a smaller fraction (2.02%) of the total threat to that species can be attributed to final consumers in the US, due to the different mix of threat causes to *Atelopus* and the different mix of implicated products consumed in the USA. These impact maps are summed over all species so hotspots, so in the Fig. 1 example the hotspot at the intersection of *Ateles* and *Atelopus* is shaded at the 2.43% + 2.02% = 4.45% level. The final biodiversity footprint map for a given country is thus a product both of the actual distribution of biodiversity hotspots around the world and the unique composition of how that country’s consumption affects each individual species in each partner country.

**Fig. 1.**
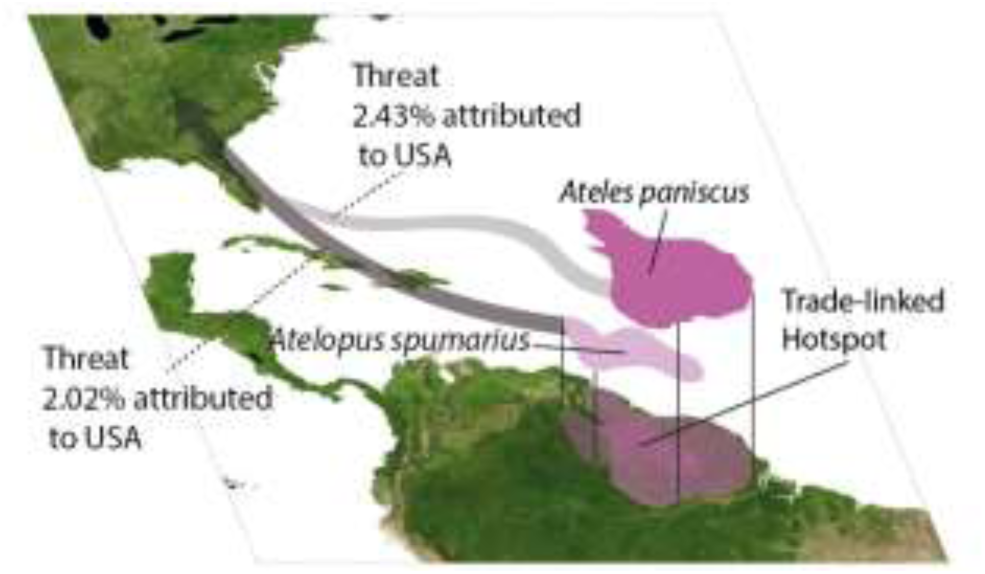
Protocol illustration, for hotspot induced by consumption in the USA.

The biodiversity footprint of country *s*, 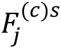 comprising the sum of the threat to species suffered in country *r* exerted directly by industry *i* due to consumption in country *s* of the good or service *j*, inclusive of the upstream and indirect impacts involved in provisioning *j* can be expressed as

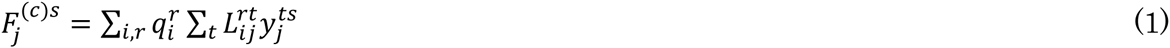

where *q* is a threats coefficients, *L* is the Leontief inverse, and *y* is final demand^1,9^. This trade model follows flows through multiple trade and transformation steps, even via middleman countries, to attribute impacts from production in *s* to consumption in *r* via last supplying country *t*. In this study we used the implementation of the biodiversity footprint from Lenzen *et al*. directly, which in turn uses the Eora global MRIO^10^. The reader is referred to that paper and its supplementary information for a thorough discussion of the method, but we summarize it briefly here.

We extend the biodiversity footprint method previously produced by Lenzen and colleagues with spatial data. Following that method we considered only species which the IUCN and BirdLife International list as vulnerable, endangered, or critically endangered, and we ignore threats which are not directly attributable to legal economic activities, including disease, invasive species, fires, and illegal harvesting (since illegal activities are not captured in the global trade model). The IUCN documents 197 different threats, 166 of which can be attributed to human activities. Threatened species hotspots were identified by overlaying species range maps from IUCN ^5^ and BirdLife International^6^ for *N* = 6,803 *Animalia* species (the combined IUCN and BirdLife databases report on 20,856 species with known threat causes, of these 8,026 are threatened, and of these range maps are available for 6,803). Species threat records from the IUCN Red List of Threatened Species, e.g. “The vulnerable (VU) *Atelopus spumarius* in Brazil is threatened by Logging and Wood Harvesting” are mapped to economic production sectors – in this case, attributed to the forestry sector in Brazil – and the resultant products are then traced through a multi-region input-output table which documents the trade and transformation steps in the economic network consisting of 14,839 sectors/consumption categories across 187 countries. When a species faces multiple threats all threats are given equal weight, as no relevant superior data is available. Every individual species is given equal weight, regardless of its ecological niche or threat level (vulnerable/endangered/critically endangered), and every threat cause is given equal weight.

The hotspot maps are potentially overestimates, for several reasons. Global MRIO databases do not currently trace flows at the sub-national level, i.e. which cities produce or consumer which goods. However the IUCN threat maps document the mix of species-threats occurring in each grid cell, and the trade model links the threats to implicated industries and traces the mix of goods and services embodied into supply chains bound for domestic or foreign final consumption. Multi-scale MRIOs combining inter-national and sub-national flows that would offer further improvements in resolution accuracy are under development ^11–14^. Another reason that the footprint maps are overestimates is that for species whose range spans multiple countries, there is no data on whether the threat(s) faced by that species occur differently in the various countries. Mathematically, our model treats country-species-threat tuples as the unique item, while in fact in the Red List it is only the [country-species] and [species-threat] tuples that are unique. For example a species spanning two countries could be threatened by logging, but it could be that logging practices in one of the two countries do not actually threaten the species, or that one of the countries does not even have any logging industry. However in the latter case, that one country has no logging industry or exports to the focal country, the range of the species in the innocent country will not be shaded since the shading is a function of the unique mix of species and export of implicated goods.

While the hotspot areas identified in this study are potentially over-estimated, it is important to note that the entire analysis is based on historical records of species threats, not current or emerging threats. Threats such as invasive species, illegal activities, or disease can arise very quickly and the Red List and Eora MRIO database could be slow to identify these current issues. This delay is particularly relevant given recent indications that humanity is surpassing “safe” limits for biodiversity loss.^15^

In this study only terrestrial and near-shore marine biodiversity are considered. Open ocean fishing was deemed beyond the scope for this study because there are a number of challenges related to getting reliable data on deep-sea fishing, both on production (handling illegal and under-reported catch), and on correctly allocating catch to the producing country (foreign-flagged vessels). Additionally, instead of the IUCN EOO maps for marine species it could be preferable to use spatialized species density models^16^ which could provide more accurate marine biodiversity hotspots. We will note that marine biodiversity is higher in coasts than open oceans^17,18^, and that jurisprudence only holds within EEZs, these two facts partially justifying the omission of extra-EEZ threats.

In this study, we link a hotspot EOO map *R* (which for display we have rasterized to 0.94′, or ~3km^2^ grid cells at the equator, though as discussed the actual accuracy of the map is less) and biodiversity footprint for each threatened species *h* for each country. Most threats (roughly 2/3rds) are exerted domestically so the country of export and the country of the hotspots are the same^1^. If a species is threatened by climate change (*CC*), the driver (exporting) country, *r*, and the suffering country, *u*, are different. Therefore, we attribute the threat to all industries who emit carbon dioxide emissions but keep the species range map in suffering countries. The unit of the resulting maps is number of species, also called species-equivalents. This value can be fractional, since one species can be threatened by many industries and countries. The footprint maps are defined as

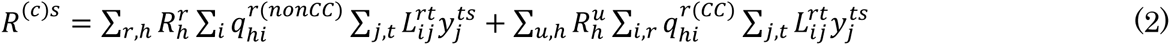

## Results

With the complete spatial footprint accounts in hand we may ask which countries, and which consumption categories, threaten habitat at various hotspots. Fig. 2 presents the biodiversity threats map driven by US consumption.

**Fig. 2.**
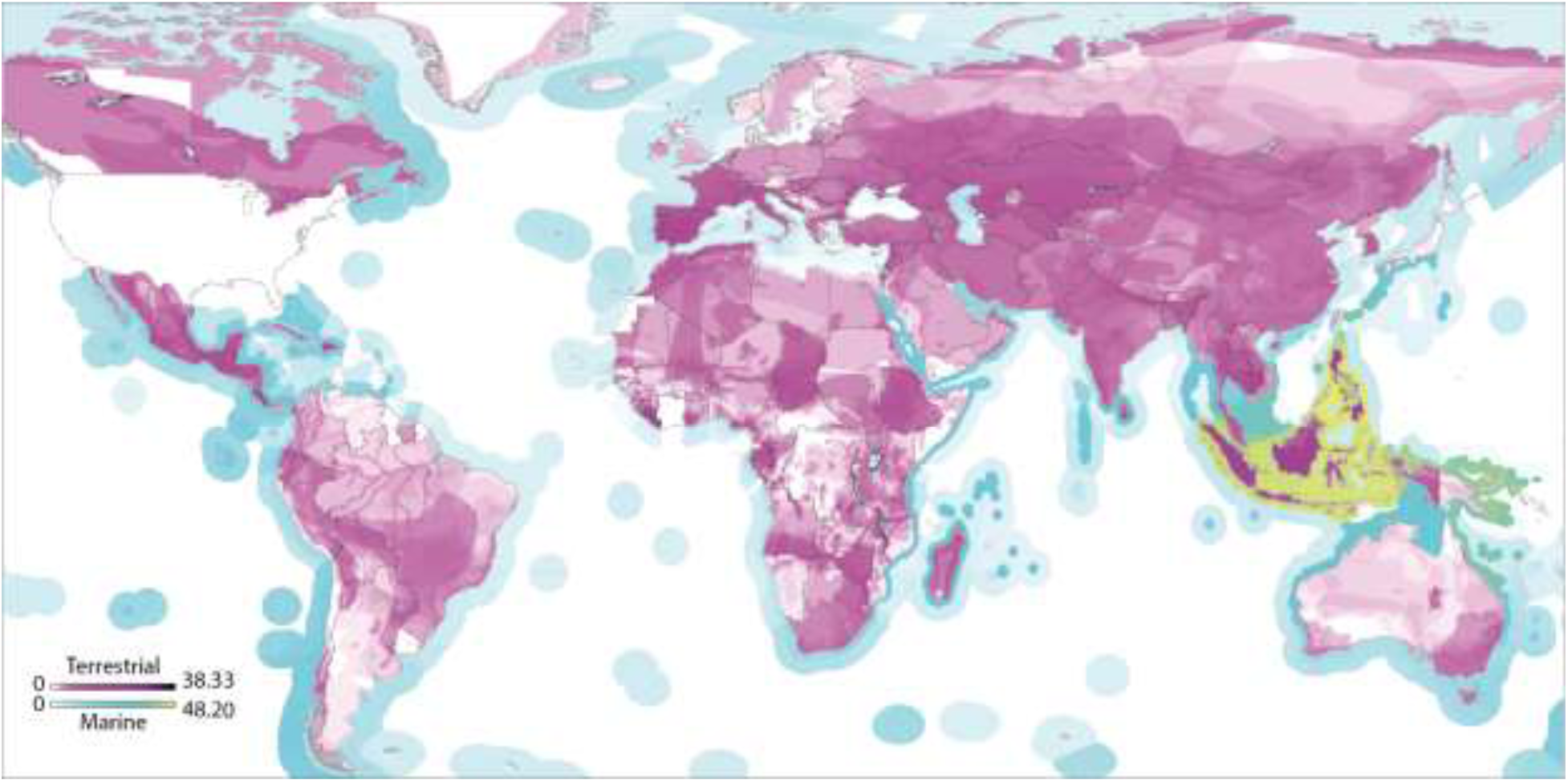
Global species threat hotspots linked to consumption in the USA. Darker areas indicate areas of threat hotspots driven by US consumption, based on the mix of threats exerted in each country and the mix of export goods sent to US for final consumption. Units are species-equivalents and are fractional since one species threat can be attribute to multiple consumer countries..

For marine species Southeast Asia is the overwhelmingly dominant global hotspot area, with the US and EU both exerting many threats there, primarily due to fishing, pollution, and aquaculture. The US has additional marine hotspots off the Caribbean coast of Costa Rica and Nicaragua at the mouth of the Orinoco around Trinidad and Tobago (Fig. 3a). The EU drives threats hotspots outside Southeast Asia in the islands around Madagascar: Réunion, Mauritius, Seychelles.

**Fig. 3.**
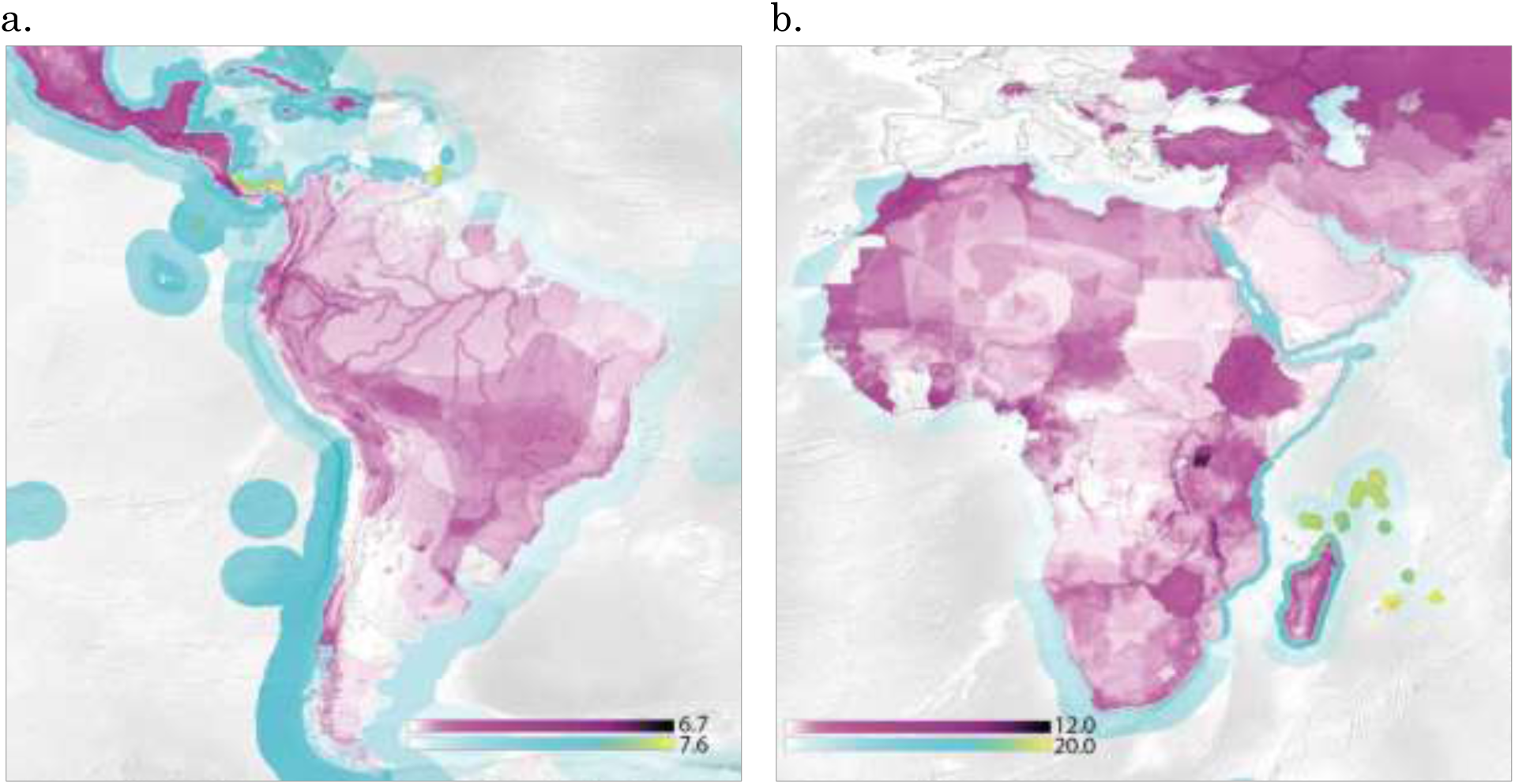
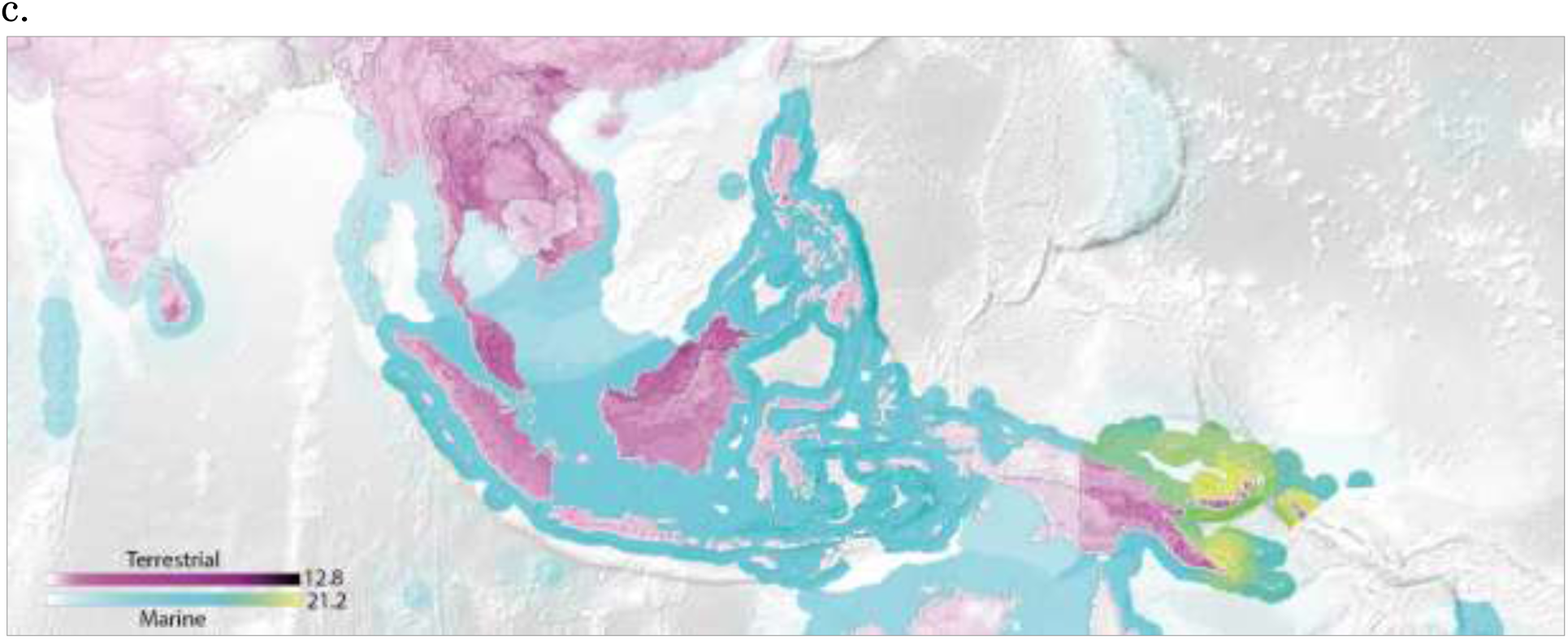
Selected enlargements of threat hotspots in Latin America driven by US consumption (panel a), in Africa driven by European (EU27) consumption (panel b), and in Asia driven by Japanese consumption (panel c). Note some countries (incl. Solomon Islands, Guyana, French Guiana, Equatorial Guinea, Western Sahara) are not covered in the economic database. Complete images are available in the Supplementary Information.

The US footprint on terrestrial species provides some notable findings. While the hotspots in Southeast Asia and Madagascar are perhaps expected, we also observe hotspots in southern Europe, the Sahel, the east and west coast of southern Mexico, throughout Central America, and Central Asia and southern Canada. Despite much attention on the Amazon rainforest, the US footprint in Brazil is actually greater in southern Brazil, in the Brazilian Highlands where agriculture and grazing are extensive, than inside the Amazon basin, although impacts along the Amazon river itself are high. The high US biodiversity footprint in southern Spain and Portugal – linked to impacts on a number of threatened fish and bird species – is also noteworthy given that these countries are rarely perceived as threat hotspots.

We find that the biodiversity footprint is concentrated: for threats driven by US consumption, the 5% most intensively affected land area covers 23.6% of its total impact on species, and at sea the 5% most intensively impacted marine area affects 60.7% of threatened species habitats.

It is possible to view the threat hotspots for various major consumer countries and zoom in on particular regions impacted by their consumption. The enlargements-in subpanels in Fig. 3 focus on threat hotspots in South America driven by US consumption (panel a), in Africa driven by EU consumption (panel b), and in Southeast Asia driven by Japanese consumption (panel c). EU consumption drives threat hotspots in Morocco, all along the coast of the Horn of Africa from Libya to Cameroon, in Ethiopia, Madagascar, throughout Zimbabwe, and at Lake Malawi and Lake Victoria. We also note the heavy EU footprint in Turkey and Central Asia; regions perhaps not known for their charismatic species but nevertheless important areas of EU-driven biodiversity impact.

The Japanese-driven biodiversity impacts in Southeast Asia are greatest in the Bismark and Solomon Sea off Papua New Guinea, and terrestrial hotspots can be found at New Britain Island (where palm oil, cocoa, logging, and coconut plantations are the dominant industries) and the eastern highlands of New Guinea; Bornean and continental Malaysia, Brunei (where urban and industrial areas sprawl into high-value habitat), the Chao Phraya drainage of Thailand, in northern Vietnam, and around Colombo and southern Sri Lanka (where pressure is driven by tea, rubber, and threats linked to manufactured goods sent to Japan).

Biodiversity footprint hotspots are a function of both underlying species richness and the level of threatening activity (i.e. number of species threats attributable to implicated industries at a given location). Hotspot maps may be decomposed to view the contributing factors. Fig. 4 provides a disaggregation of the USA footprint in Fig. 2 by threat cause. Hotspots are a function of underlying species density (composited EOO maps).

**Fig. 4.**
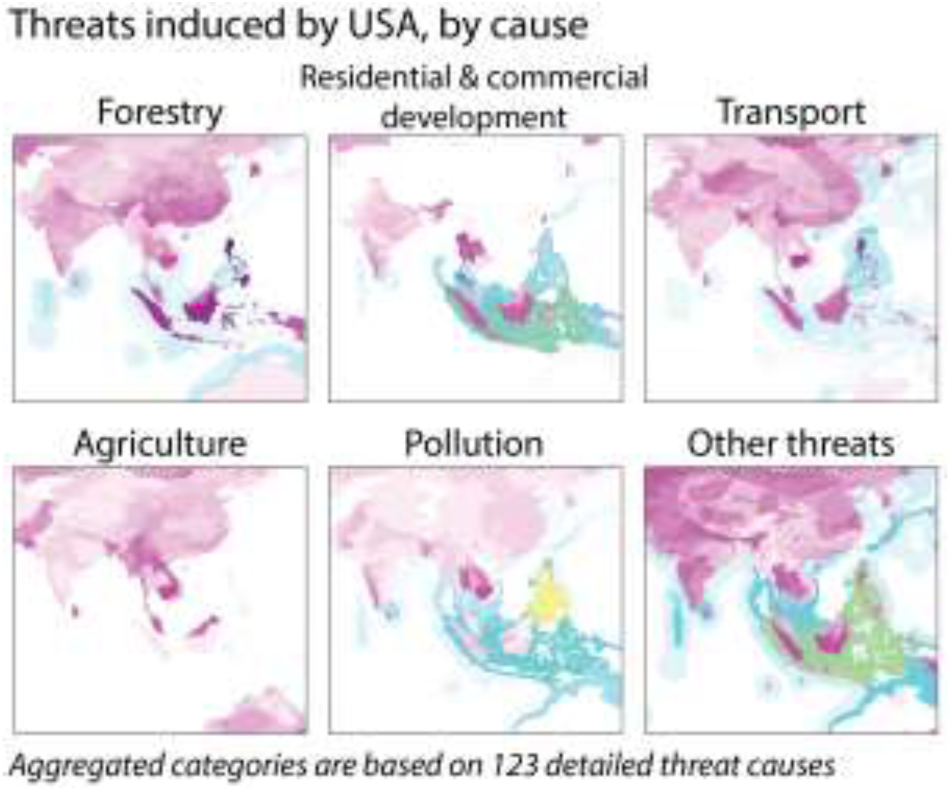
Decomposition of threat hotspots linked to consumption in the USA (see Fig. 2) by threat cause. Biodiversity footprint hotspots for a given country are a function of both underlying species richness and the composition and volume of threatening activity.

## Discussion, Limitations, and Conclusions

Trade and responsibility attribution aside, identifying biodiversity hotspots is not trivial, and is strongly limited by data resolution. The need for improved models and maps locating species occurrence and biodiversity hotspots is documented^19,20^. Since Myers and colleges introduced the hotspot concept with 25 broad areas,^21^ much conservation research now relies on extent-of-occurrence (EOO^22^) maps. Overlapping EOO maps^23^ has limitations as a method for finding hotspots^24–28^, and extent of occurrence is not the only way to identify hotspots: for birds, mapping species occupancy^29^, endemism, or threat reveals different hotspots^30^. Furthermore, both threat intensity and species density can vary considerably within the range^31^. Projects such as AquaMaps^16^, the Global Mammal Assessment^18^, and the Global Amphibian Assessment are working to generate more robust and high-resolution maps. When a superior, globally consistent, set of species occurrence maps becomes available it will be possible to replace the EOO maps with those. Grenyer and colleagues^32^ argued that priority areas for biodiversity conservation should be based on high-resolution range data from multiple taxa, not merely aggregated extent of occurrence maps since cross-taxon and rare species congruence are in fact low in such aggregate maps. Acknowledging this, the method we use here can be used to identify the spatial biodiversity footprints at the detail of individual species. It is also possible to use the spatial footprinting method with biodiversity threat hotspot maps generated using other approaches such as mechanistic modelling. Since EOO maps of range do not estimate actual occupancy or how threat varies across the range, more detailed local assessments at individual hotspots will be always be needed. Nevertheless these spatial footprint maps can be of use. For example, we can imagine that even if a company or buyer consults a spatial biodiversity footprint map that has overestimated the threat and identifies, say, three hotspots in a supplier country, even though over-estimated (i.e. the true hotspots will be in some sub-set of the identified area), this hotspot information is still more precise and actionable than simply a single total figure for impacts in that country, which has been the limit of knowledge so far.

The economic trade model is another source of uncertainty, although work continues to improve the convergence^33^, reliability, spatial^11,12^, and product-level detail^34^ of multi-region input-output databases used for the trade accounting. While alternative methods exist to calculate land footprints^35^ – which is the biggest driver of the biodiversity footprint^36^ – for this study an existing biodiversity footprint account was used rather than building a new one. Improved spatial data of the trade model is especially important for spatially extensive countries such as USA, China, Russia, and India, where one industry may have different impacts across its domain. With much attention on global supply chains^37^ and footprints^38^ it may be expected that the trade accounting and embodied resource flow accounts will become more accurate in the future. However it must be noted that small-scale and illegal impacts are potentially important^39^ and will possibly never be covered by global-scale trade databases.

It has been estimated that 90% of the $6 billion of annual conservation funding originates in and is spent within economically rich countries^40^, yet these countries are rarely where threat hotspots lie. Directing funding back up along their supply chains, toward the original points of impact, could help yield better conservation outcomes.

As conservation efforts must both protect critical habitat^41,42^ and do so in an economically efficient manner^43–45^ spatially explicit supply chain analysis can be a helpful tool for finding the most efficient ways to protect absolutely important areas. Given that the Aichi targets are inadequate –protecting 17% of global land area and 10% of marine can cover at most 53.1% of known threatened species EOO – and the fact that biodiversity stocks are not evenly distributed amongst countries, conservation hotspots must be prioritized (and of course, not be placed in unproductive areas^46–48^).

Shortly before this paper came to print we became aware of another spatially explicit biodiversity footprint study by Kitzes and colleagues that used estimated species richness based on potential net primary productivity to estimate which economic activities impact the most potentially valuable bird habitats.^49^

Using the biodiversity hotspots method it is possible to identify areas where the biodiversity threat is predominantly driven by just a small number of countries. By identifying regions where just two or three countries are implicated in driving the pressure it could be easier to initiate direct collaborations between producers and consumers, in parallel to existing international regimes, to mitigate biodiversity impacts at those places.

Spatially explicit impact accounting can facilitate improvements in sustainable production, international trade, and consumption. Responsibility for environmental pressures should be shared along the supply chains, not pinned solely on primary impacting industries nor exclusively on final consumers. Looking upstream, detailed information on species hotspots can be useful for companies in reducing their biodiversity impact. Downstream, accounts such as these can be of use to guide sustainable purchasing and green labelling and certification initiatives. It is possible to imagine companies comparing maps of biodiversity footprints with maps of where their inputs are sourced. We could also foresee conservationists working to preserve impacted areas using such models to help identify the intermediate and final consumers whose purchases sustain threat-implicated industries, and looking down the supply chain to help involve consumers in protection activities. Better targeting spatial hotspots assist in setting effective conservation priorities^50^.

Maps of species threat hotspots can help all actors, from producers and conservationists to final consumers, to focus solutions on targeted biodiversity hotspots.

## Acknowledgements

This work was supported in part by the Japan Society for the Promotion of Science through its Grant-in-Aid for Young Scientists (A) 15H05341, the Norwegian Research Council grant #255483/E50, the PRINCE project of the Swedish Environment Agency, and Belmont Forum’s TSUNAGARI project. The authors would also like to thank A. Hart for comments that have improved the work.

## Author contributions

Both authors contributed equally to this work.

## Data Availability

The results maps presented and discussed in this paper are available in the online Supplementary Information package. The results, calculated as described in the Methods, are based on the data from the IUCN, BirdLife International, and the Eora MRIO databases, all of which are publicly available.

The authors affirm they have no competing interests.

